# No Replication of Direct Neuronal Activity-related (DIANA) fMRI in Anesthetized Mice

**DOI:** 10.1101/2023.05.26.542419

**Authors:** Sang-Han Choi, Geun Ho Im, Sangcheon Choi, Xin Yu, Peter A. Bandettini, Ravi S. Menon, Seong-Gi Kim

**Affiliations:** Center for Neuroscience Imaging Research, Institute for Basic Science, Suwon, Republic of Korea; Athinoula A. Martinos Center for Biomedical Imaging, Department of Radiology, Harvard Medical School, Massachusetts General Hospital, Charlestown, Massachusetts, USA; Section on Functional Imaging Methods and Functional MRI Facility, NIMH, NIH, Bethesda, MD, USA; Centre for Functional and Metabolic Mapping, Robarts Research Institute, Western University, London, Ontario N6A 5B7, Canada; Department of Biomedical Engineering, Sungkyunkwan University, Suwon, Republic of Korea

**Keywords:** direct neuronal activity, fMRI, sample size, noise distribution

## Abstract

Direct imaging of neuronal activity (DIANA) by fMRI could be a revolutionary approach for advancing systems neuroscience research. To independently replicate this observation, we performed fMRI experiments in anesthetized mice. The BOLD response to whisker stimulation was reliably detected in the primary barrel cortex before and after DIANA experiments; however, no direct neuronal activity-like fMRI peak was observed in data of individual animals with the 50-300 trials. Extensively averaged data involving 1,050 trials in 6 mice showed a flat baseline and no detectable neuronal activity-like fMRI peak. However, spurious, non-replicable peaks were found when using a small number of trials, and artifactual peaks were detected when some outlier-like trials were excluded. Further, no detectable DIANA peak was observed in the BOLD-responding thalamus from the selected trials with the neuronal activity-like reference function in the barrel cortex. Thus, we were unable to replicate the previously reported results without data pre-selection.

## Introduction

Blood oxygenation level-dependent (BOLD) functional magnetic resonance imaging (fMRI) has revolutionized how neuroscientists investigate human brain functions and networks. (*1–3*). However, BOLD fMRI measures hemodynamic responses as a surrogate of neuronal activity, thus spatial and temporal resolution is highly dependent on BOLD sensitivity and neurovascular coupling (*4*). To further understand brain functions at the mesoscale, it would be ideal to measure causality and sequences of neuronal events by fMRI. Initial approaches have been to utilize differences of BOLD fMRI onset times between layers or regions (*5–9*). The rationale for our ultra-high resolution, capillary-specific studies was that since the contribution of capillaries to BOLD fMRI increases with magnetic field strength, and these capillaries are in close proximity to neurons, early hemodynamic responses at ultrahigh fields could reflect changes in neuronal activity with greater fidelity. However, while this approach is feasible in ultrahigh fields with extensive averaging, it may be beyond the capabilities of current human MRI systems.

Recently, Toi et al. reported that direct imaging of neuronal activity (DIANA) can be achieved by high temporal resolution fMRI in the anesthetized mouse model at 9.4 T (*10*). When a whisker pad electric stimulus was applied, the peak intensity of ∼0.15% was detected at ∼12 ms in the thalamus and ∼25 ms in the primary somatosensory cortex following the stimulus pulse. When optogenetic stimulation was applied in the primary sensory cortex (S1), the peak intensity of ∼0.3% was observed at ∼15 ms in the S1 and ∼25 ms at the thalamus after the stimulus onset. These DIANA data were shown to have low variability among animals and were highly consistent with electrophysiology latency data. Although low temporal signal-to-noise ratio (tSNR) is expected due to high spatial (220×220×1000 µm^3^) and temporal resolution (5 ms) in conjunction with an intrinsically noisy acquisition method, 10.8-s long trials were averaged only 40 times for whisker stimulation and even less (13-20 times) for optogenetic stimulation. This breakthrough fMRI approach has an unusual combination of high spatial resolution, high temporal resolution, high detectability, low inter-animal variation, and most importantly, apparent direct neuronal detection.

Many laboratories around the world have been attempting to reproduce DIANA signals in various experimental conditions without success. In preliminary human studies (*11*), DIANA signals were not observed, possibly due to species difference (human vs. mouse) or subject status (awake vs. anesthetized). Although data in the original publication (*10*) appear convincing under multiple experimental manipulations, it is critical to independently reproduce these DIANA findings. Thus, we repeated the reported whisker stimulation experiments in virtually the same mouse model used in Toi et al. (*10*). Compared to the original study, two improvements were made; 1) continuous infusion of anesthetics (*12, 13*) rather than intermittent bolus injection of anesthetics (*9, 14*) for maintaining stable animal physiology during fMRI studies, and 2) the use of 15.2 T rather than 9.4 T for enhancing SNR (*15*). All other parameters were the same as in Toi et al. (*10*).

## Results

Six anesthetized mice were used for fMRI studies of whisker pad stimulation at 15.2 T (Fig. 1A). For anesthesia, a mixture of ketamine and xylazine was initially injected intraperitoneally (IP) and was continuously infused intravenously (*12, 13*) in order to maintain stable animal physiology. Toi et al. (*10*) used the bolus IP delivery of supplementary anesthetics when needed (*14*), which results in well-known modulation of anesthetic depth during experiments (*14, 16*). For whisker stimulation, electrodes were placed on the mouse’s right whisker pad (*12*). Stimulation parameters included a pulse width of 0.5 ms and current intensity of 0.5 mA.

**Figure 1.**
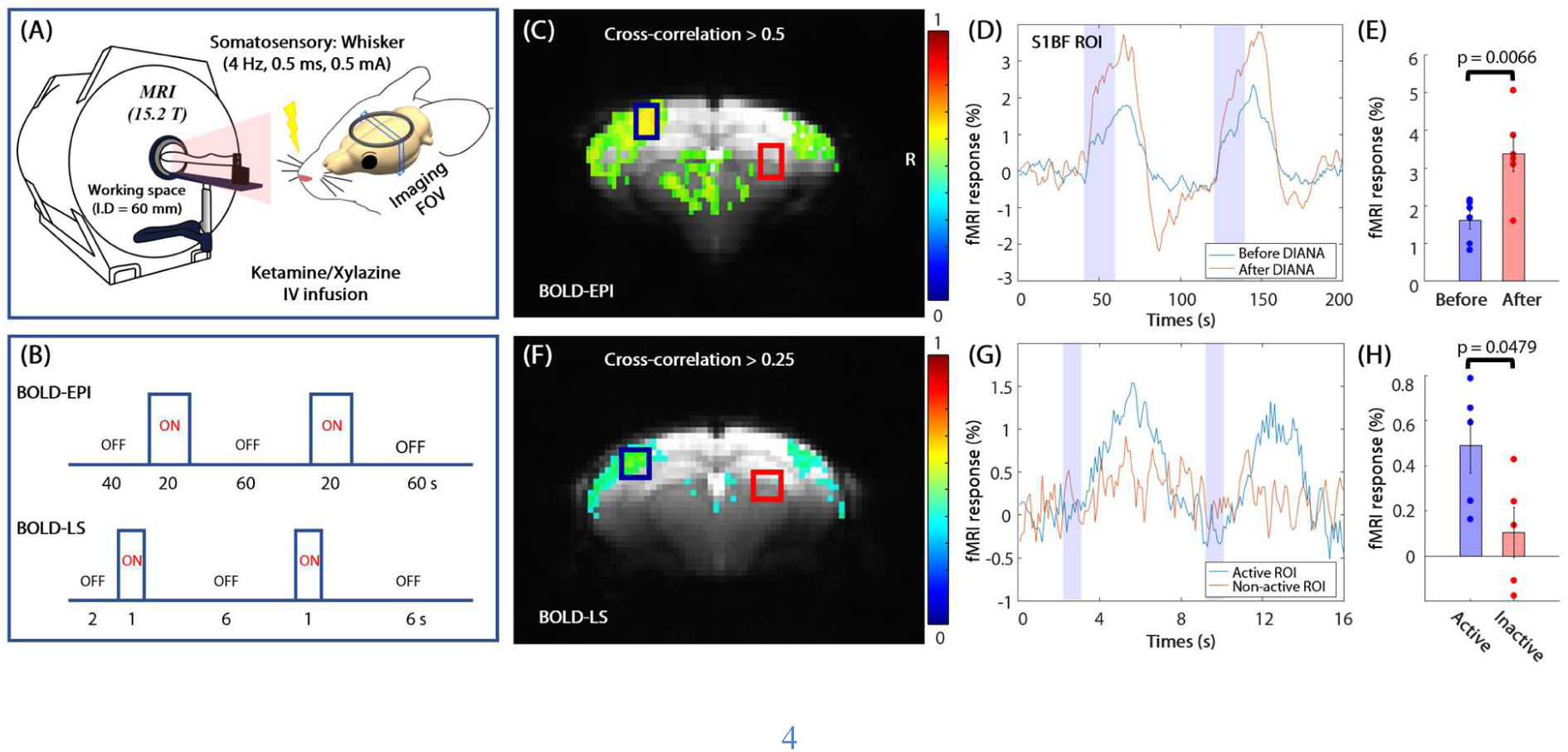
BOLD fMRI of somatosensory stimulation in anesthetized mice at 15.2 T. (**A** and **B**) Experimental schematics and stimulation paradigms of BOLD fMRI with EPI and line scanning (LS). (**C**) BOLD fMRI map overlaid on original EPI image of one animal (Mouse #1). Black box: active 5×5-voxel S1BF ROI; red box: inactive 5×5-voxel ROI. (**D**) BOLD time courses of the active S1BF ROI before and after DIANA experiments in Fig. 1C. Vertical shaded bars: stimulus durations. (**E**) Average percent changes of the S1BF ROI before and after DIANA experiments (n = 6 mice). Each data point: each mouse. (**F**) Conventional BOLD fMRI map obtained from only one shuffled line scanning fMRI run. (**G**) Time courses of active and inactive ROIs in Fig. 1F. Vertical shaded bars: stimulus durations. (**H**) Averaged percent changes of active and inactive ROIs (n = 5). Each data point: each mouse; error bars: SEM.

### BOLD fMRI responding to whisker stimulation at 15.2T

All fMRI data were acquired at 15.2 T for enhanced sensitivity (*15*). One 1-mm thick slice covering the primary somatosensory barrel field (S1BF) was chosen using scout BOLD fMRI studies to identify S1BF. To ensure that anesthetized animal’s condition enabled reliable detection of the stimulus over the duration of the imaging session, BOLD fMRI studies were performed using standard gradient-echo (GE) echo planar imaging (EPI) with repetition time (TR) of 1 s and echo time (TE) of 11.5 ms before and after DIANA experiments (Fig. 1B). These control experiments to confirm stable neuronal activity over the entire imaging session utilized 200-s runs with two blocks of 20-s whisker stimulation. One animal’s fMRI data are presented in Fig. 1C-D. The BOLD map responding to whisker pad stimulation at 4 Hz (Fig. 1C for one mouse) shows reliable activation in the contralateral S1BF area and contralateral thalamus. Two 5×5-voxel regions of interest (ROI) were chosen for further time course analyses, an ROI in the activated area of the contralateral S1BF (“active ROI”) and an ROI in the inactive ipsilateral thalamus (“inactive ROI”). The post-DIANA BOLD fMRI response in the active ROI was higher than the earlier pre-DIANA BOLD response (Fig. 1D for one animal’s time courses, and Fig. 1E for individual data), which is consistent with our previous time-dependent BOLD fMRI studies (*12*). This indicates that BOLD fMRI responses are intact pre- and post-DIANA, confirming the presence of stable neuronal activity. Thus, the animals’ physiological state should allow for the detection of a stimulus in the DIANA experiments.

To ensure that the DIANA acquisition method, a shuffled line scanning k-t pulse sequence originally developed by Silva and Koretsky (*6*), was working correctly, we performed BOLD fMRI with TR/TE of 100/11.7 ms, flip angle = 17° (Ernst angle) in the S1BF, and in-plane resolution of 0.22 × 0.22 mm^2^. Acquisition of 160 frames was repeated for 54 phase-encoding steps, resulting in experimental run time of 16 s × 54 = 864 s. The BOLD response to 1-s whisker stimulation was reliably detected - even from one run of 54 stimulus events (Fig. 1F-G). An average signal change in the contralateral S1BF area was 0.49 ± 0.27% (SD, n = 5), while small or negligible response was observed in the ipsilateral thalamus ROI (Fig. 1G-H). Our data indicate that the shuffled k-t pulse sequence was implemented correctly. In this pulse sequence, each subsequent line of k-space that forms a given image is separated in time by 16 s, so the shuffled k-t approach is highly sensitive to physiological noise and instrument instability and is thus intrinsically more noisy despite 1 s × 54 phase-encoding steps = 54 s of total stimulation for a single BOLD-LS run versus 20 s × 2 runs = 40 s in BOLD-EPI, leading to lower correlation and p values for the k-t approach compared to EPI.

### DIANA fMRI with high temporal resolution at 15.2T: Sensitivity and number of averages

Once we validated the k-t pulse sequence with BOLD, in order to reproduce DIANA findings in anesthetized mice with the same imaging parameters from Toi *et al.* (*10*), we used the shuffled line scanning k-t pulse sequence with TR/TE of 5/2 ms and spatial resolution of 0.22 × 0.22 × 1 mm^3^. According to the postulated mechanism of the DIANA effect (*10*), DIANA responses at 15.2 T should be enhanced due to larger T_2_ weighting compared to 9.4 T. This is because the T_2_ of tissue water at 15.2 T is shorter than that at 9.4 T (24.5 vs. 35.7 ms) (*17*) and DIANA peaks are linearly dependent on TE (*10*). Acquisition of 40 (200 ms) or 200 frames (1000 ms) was repeated at 54 phase-encoding steps, lasting 10.8 s or 54 s for each run. In each mouse, at least 50 runs were repeated for each experimental condition (see table S1 for experimental design). Since an average of only 40 10.8-s trials at 9.4 T showed a 0.17% peak response in the S1BF ROI at 25 ms after the stimulus in the original publication (see Fig. 1D in (*10*)), we expected to easily detect the DIANA peak with 50 averages at 15.2 T given that tSNR of thermal-noise-dominant high-resolution fMRI increases with magnetic field strength (*15, 18, 19*).

Since 40 trials were used in the original DIANA paper (*10*), 50 trials with 1000-ms duration (200 frames) were analyzed in detail (Mouse #3). tSNR was calculated by the signal mean divided by signal standard deviation over the 200 frames. Within the S1BF ROI, average voxel-wise tSNR was 32 for a single trial (ranging between 28 and 42 in 6 animals), and 230 for 50 trials-averaged data (Fig. 2A). The activation map obtained using a voxel-wise cross-correlation analysis with the DIANA response function (*10*) did not show any visible activation cluster in the S1BF (Fig. 2B). Similar observations were detected in all 6 mice. This may be explained by the low sensitivity of the direct neuronal-related fMRI response on a single voxel level.

**Figure 2.**
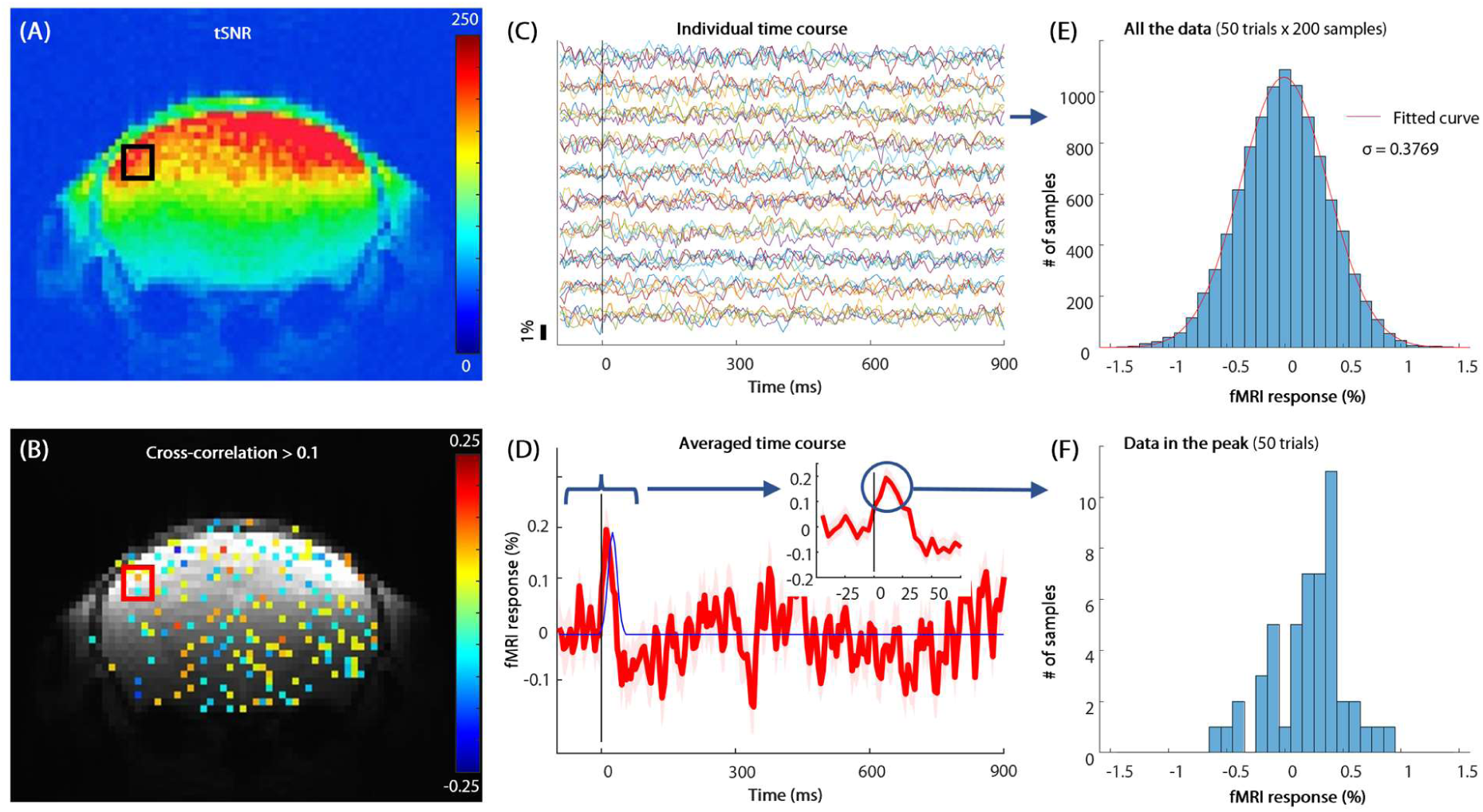
Systematic analysis of 50 DIANA trials in one animal. One DIANA trial composed of total 20 pre-stimulus and 180 post-stimulus frames (i.e., 1000-ms inter-stimulus interval) (Mouse #3, see table S1). (**A**) tSNR map calculated from the average of 50 trials. (**B**) Cross-correlation map obtained with an expected neuronal response curve from Toi et al. (*10*), which was thresholded by absolute cross-correlation values of 0.1. Square box: 5×5-voxel S1BF ROI. (**C**) Trial-wise time courses of the active S1BF ROI. To visualize individual trials, 5 trials were plotted per row. Vertical bar: stimulus. (**D**) The averaged time course of 50 repeated trials with an expanded view in inset (red). An expected DIANA response (peaked at 25 ms) based on electrophysiology was also plotted (blue). A statistically significant positive peak was detected at 10 ms after the stimulus. Shaded area: SEM. (**E**) A histogram of Figure 2C data points (50 trials × 200 time points). A Gaussian noise distribution was observed with a standard deviation of 0.377%. (**F**) A histogram of 50 individual trials at a fixed time (10 ms post-stimulus) of the statistically significant positive peak. DIANA-like peaks may be observed from noisy data with limited samples just by chance.

To increase tSNR and detectability, time courses were obtained from 5×5-voxel ROIs determined from the scout BOLD experiments as described, using a moving average of 3 frames (Fig. 2C). Assuming that all data points are independent, this form of spatial averaging and temporal smoothing increases tSNR by (number of voxels × number of temporal averages)^1/2^. As expected, the measured tSNR increases with temporal averages (first 10, 20, 30, 40, and 50 trials in fig. S1) and the number of number of voxels (1, 9 and 25 voxels in fig. S1). Trial-wise tSNR of 5×5-voxel S1BF ROI across 200 frames was 279.1±23.5 (SD, n = 50 trials), and tSNR of the all-averaged time series was 1585 for the entire 200 frames and 2628 for the initial 20 frames. If stimulus-related responses were significant, tSNR obtained from the entire time series of frames would in fact be underestimated.

The direct neuronal activity-related fMRI peak of 0.17% in the original DIANA paper (*10*) can be easily detected from the 50 trials-averaged 15.2 T data with tSNR of >1500. The averaged time course (Fig. 2D) showed a direct neuronal activity-like peak of 0.2% in the S1BF ROI right after somatosensory stimulation and another peak of −0.15% at 340 ms. When pre-stimulus data points (20 frames × 50 trials) were compared with data in the peak, both peaks were statistically significant (p = 8.7e-05 and 0.0025, two-sample t-test). However, the positive peak latency of 10 ms (Fig. 2D inset time course) did not match with the neuronal time to peak of 25 ms (*10*), thus our level of certainty that this peak reflects neuronal activity is low.

The above-measured peak may be a consequence of noise in a limited sample size. When all time course data points (50 trials × 200 frames) were considered, a Gaussian noise distribution was observed (Fig. 2E). However, when only 50 trials of the peak frame (10 ms post-stimulus) were considered, a skewed distribution was observed in a histogram (Fig. 2F), indicating that a noise peak can appear statistically significant just by chance due to an insufficient, biased sample. If an observed peak is genuine, then it should be reproducible across animals and be differentiated better from the noise level when more averaging is performed. Thus, an increased number of repeated trials (see Fig. 3C, animal #3) is needed to determine whether a genuine neuronal activity-related peak can be detected.

**Figure 3.**
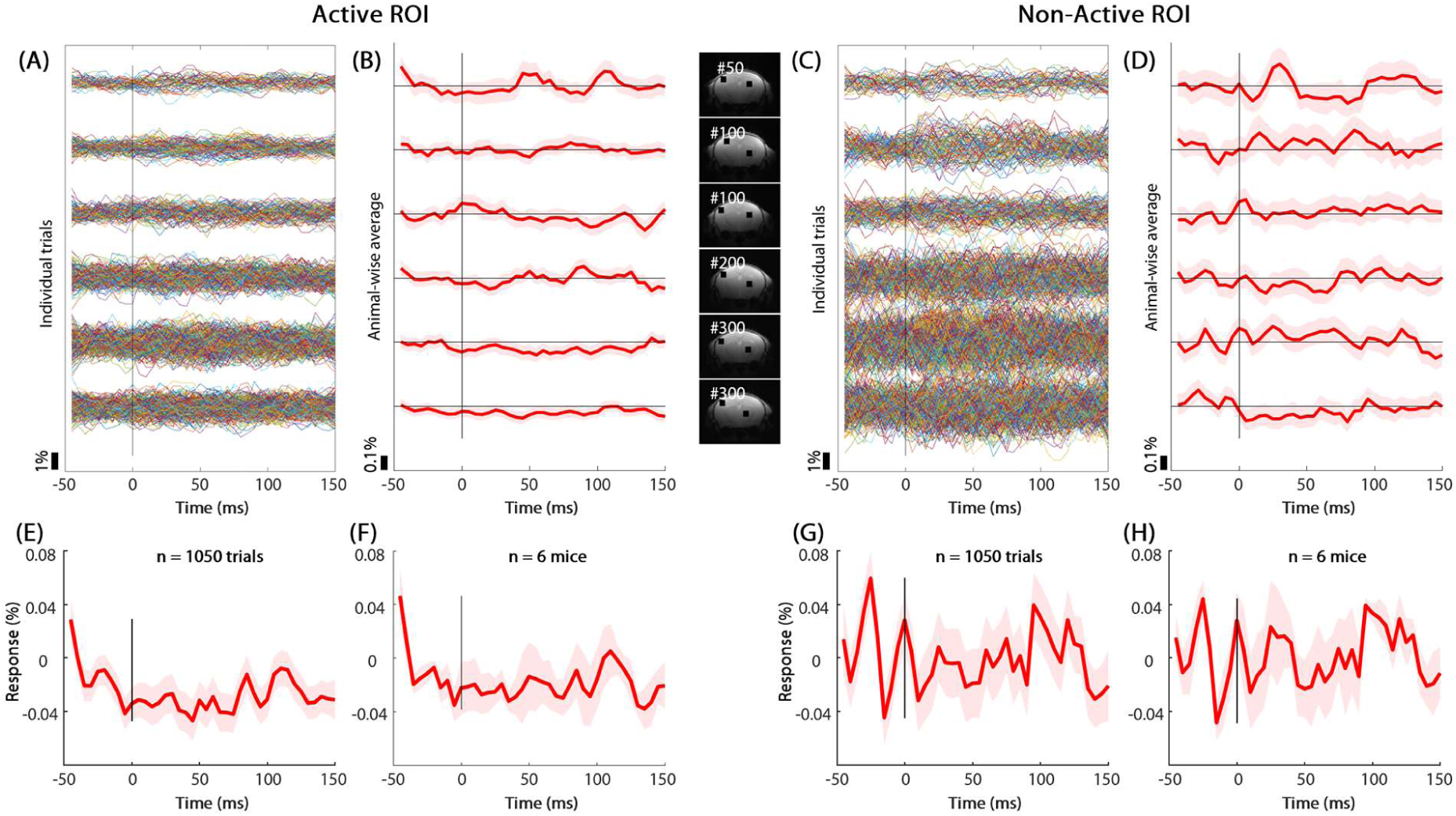
No observable DIANA responses in all six subjects. (**A** to **D**) Individual (A, C) and averaged time courses (B, D) from the active contralateral S1BF (A-B) and inactive ipsilateral ROIs (C-D) in each individual animal (row). ROIs were shown with the total number of trials in each animal (images between B and C). 50 – 300 trials were repeated in each animal. (**E** to **H**) Averages of all 1050 trials in 6 animals (E, G) and 6 averaged animal’s time courses (F, H) from the active S1BF (E, F) and inactive ROI (G, H). Note that the peak-like response at −50 ms (E-F) is the initial point of our measurements before reaching a steady-state condition. No noticeable neuronal activity-like peak was observed. Vertical bar: stimulus; shaded area: SEM.

### Extensively averaged DIANA fMRI data: No observation of direct neuronal activity-related peak

All repeated runs in each mouse were stimulation-pulse-locked, leading to 50 – 300 trials in each animal (table S1). There was no difference of baseline variations between runs with and without radio frequency (RF) spoiling. In each subject, time courses of all trials (50 – 300) were obtained from the contralateral S1BF (Fig. 3A) and inactive ipsilateral thalamic ROIs (Fig. 3C). Then, an averaged time course was obtained by averaging all repeated trials in each subject (Fig. 3B and D).

ROI-wise tSNR for individual animals with 50 – 300 averages ranged between 1950 and 4583 in the active S1BF ROI and between 1500 and 2731 in the inactive ROI. Even in averaged time series with high tSNR values, no DIANA-like activity was observed in any of the animals. Note that the statistically significant positive peak observed in Figure 2 proved to be insignificant when 100 trials were averaged (3^rd^ row in Fig. 3B). To enhance the sensitivity further, time courses were averaged from all 1050 trials in 6 mice (1050 × 54 = 56,700 pulse events) (Fig. 3E and G) and from individual animal’s averaged time courses (Fig. 3F and H). When 1050 trials were averaged, tSNR was 7370 for the S1BF ROI. Even when extensive averaging was achieved, no direct neuronal activity-like fMRI peak was detected.

### Artifactual DIANA peak can be generated by exclusion of presumably bad trials

Given our inability to detect a DIANA signal in data with extremely high tSNR and extensive averaging, we investigated how one might observe a DIANA peak when dealing with more limited sampled data. In anesthetized animal fMRI studies with the intermittent IP bolus injection of anesthetics as Toi et al. (*10*) adopted, anesthesia depth is modulated (*14, 16*). At a deep anesthesia condition (such as right after the bolus injection), neuronal activity and corresponding fMRI responses are suppressed (*14*). In such an intermittent anesthesia protocol, the observation of no BOLD response was regarded as an indication of deep anesthesia, and these runs are often appropriately excluded for further data analysis.

Throwing away bad trials is often practiced, as these trials presumably occur due to poor anesthetized animal conditions that lead to a null, independently measured BOLD response. However, it would be incorrect to use temporal similarity to the hypothesized DIANA response as a metric to select “good” from “bad” trials, as spurious noise could then average together to produce the hypothesized result. To demonstrate the impact of a biased exclusion of “presumably bad trials” on DIANA studies, we re-processed our DIANA data with 50 – 300 trials (Fig. 3). Initially, the correlation between each time series and a negative response function with a peak time of 25 ms relating to a DIANA response and 35 ms as a control (Fig. 4B) was calculated for identifying presumably outlier trials systematically. Trials were ranked based on their cross-correlation values with this function, and the top 6% (e.g., 3 out of 50 trials), 10% (e.g., 5 out of 50), and 20% ranks (e.g., 10 out of 50) were used as an outlier exclusion threshold. Then, the averaged time course of the included (94%, 90%, and 80% of all trials) or excluded trials was obtained for each subject in each exclusion threshold. The original data (Fig. 4A and fig. S2A for individual animals; Fig. 4C and fig. S2C for an average of 6 animals) show no neuronal activity-like peak. When some trials were excluded, the averaged time course of included trials for each individual animal showed noisy positive responses (Fig. 4D,E for active ROI and fig. S2D,E for inactive ROI), while the averaged time course of excluded trials showed a negative peak at a chosen time (Fig. 4D,E for active ROI and fig. S2D,E for inactive ROI). When only 10% of total trials were excluded, noisy time courses having an observable peak were similarly observed in both rejection criteria (neuronal time of 25 ms vs. control 35 ms) (Fig. 4F for active ROI and fig. S2F for inactive ROI). Peak intensity was also closely dependent on tSNR (see Fig. 4 vs. fig. S2) and the exclusion threshold; a larger spurious peak intensity was observed for a noisier dataset at a higher exclusion threshold (Fig. 4F and fig. S2F), not surprisingly as one is effectively rejecting more outlier responses and selectively averaging positive noise peaks that occur when the hypothesized signal is occurring. When data exclusion is performed based on any pre-selection model of the expected response implicitly (visual inspection) or explicitly (cross-correlation analysis), then statistical circularity occurs (20), invalidating results, as seen in Fig. 4 and fig. S2.

**Figure 4.**
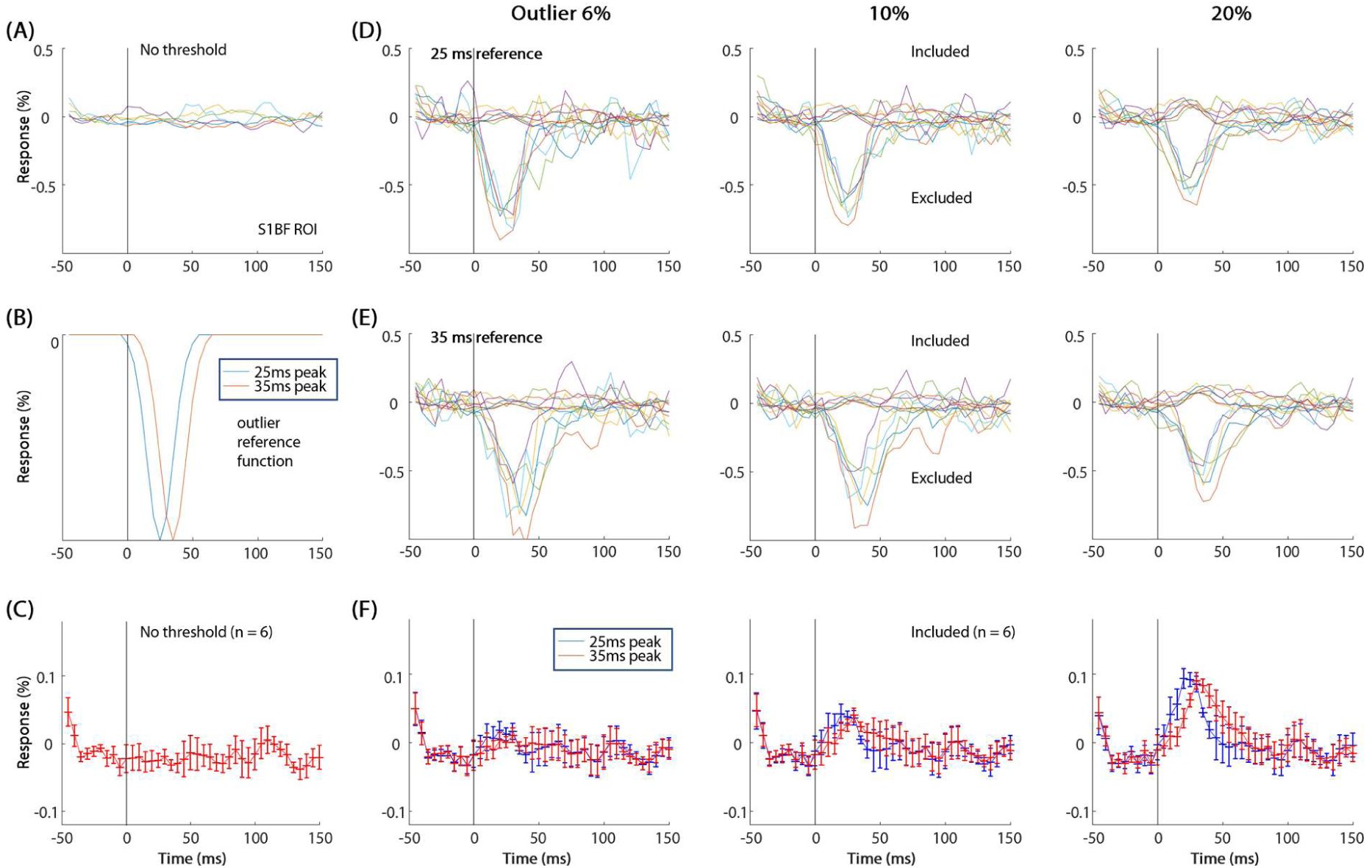
Artifactual fMRI peak induced by temporal response-based data exclusion. (**A**) Averaged time courses of six individual animals in the active S1BF ROI without excluding trials (same as Fig. 3B). (**B**) A Gaussian reference outlier function with a full width of half minimum (FWHM) of ∼25 ms peaked at 25 ms (blue) and 35 ms as a control (red). (**C**) An average of all animals’ time courses without excluding trials. This is the same as Fig. 3F. In six animals, an individual trial’s time course was correlated with a reference function shown in B, and ranked among all trials, based on its cross-correlation value. Then, in each animal, trials were separated into included and excluded categories, based on an outlier threshold of top 6%, 10%, and 20% cross-correlation values. (**D** and **E**) Averaged time courses of the included and excluded trials in individual animals for the 25 ms (**D**) and 35 ms peak reference function (**E**). Averaged time courses of the included trials showed noisy positive responses, while those of the excluded trials showed a negative peak around the expected peak time. Each color time course: each animal. (**F**) Means of subject-wise included trials time courses with the 25 ms (blue) and 35 ms peak reference function (red). Clearly, an exclusion process leads to a spurious peak from noisy data. It is fundamentally important to note that these peaks are erroneous because such a preselection process is statistically circular, summing noise in a biased manner to produce precisely the peak that was preselected. Error bars: SEM.

### Genuine neuronal activity-related DIANA peak is not observable even from highly selected trials

Despite being statistically invalid, one may think that it is acceptable to select DIANA-appearing trials if stimulation-evoked neuronal activity directly contributes to fMRI signals. If this hypothesis is valid, then the selected trials with DIANA-like responses in one region (e.g., S1BF at 25 ms peak time) should produce a DIANA peak in the anatomically and functionally networked regions (e.g., thalamus at 15 ms), resembling the findings of the thalamus and S1BF in the original paper (Fig. 2 in Toi et al. (*10*)). To evaluate this possibility, we re-analyzed time course data of 3 mice with robust BOLD-EPI responses in the contralateral thalamus and S1BF before and after DIANA experiments (Fig. 1C for fMRI map and 5A for ROIs). Each time course was correlated with an expected neuronal response template with a peak time of 15 ms in the thalamus and 25 ms in the SIBF (Fig. 5B). Trials were separately selected for S1BF and thalamus with a threshold of top 20%, 50%, and 80% cross-correlation values. Among 450 trials in three mice, the number of trials jointly selected for *both* thalamus and S1BF was 17, 110, and 288 for 90, 225 and 360 selected trials in each ROI, respectively, which can be explained by the probability of noise distributions (i.e., 450 times 0.2^2^, 0.5^2^, and 0.8^2^). Then, two different sets of selected trials for each threshold were used to obtain time courses in the S1BF and thalamus (Fig. 5B-C). Obviously, the DIANA-like peak was detected in the *selected* ROI as a consequence of circular reasoning, but only noisy time courses were observed in the *unselected* functionally networked region. Similar observations were also detected when all 6 mice were used (fig. S3). To further substantiate our ROI observations, we computed voxel-wise cross-correlation values by averaging selected trials with the top 20% cross-correlation values for the S1BF (with a one-voxel spatial and 3-point moving average) and then comparing them to a reference function of peak time shifts between 10 and 30 ms after the stimulus onset (Fig. 5D for one mouse and fig. S4 for 3 mice). The analysis revealed a significant cluster only in the ‘selected’ S1BF, not in the thalamus. In summary, we confirm that the justification of trial selection is invalid.

**Figure 5.**
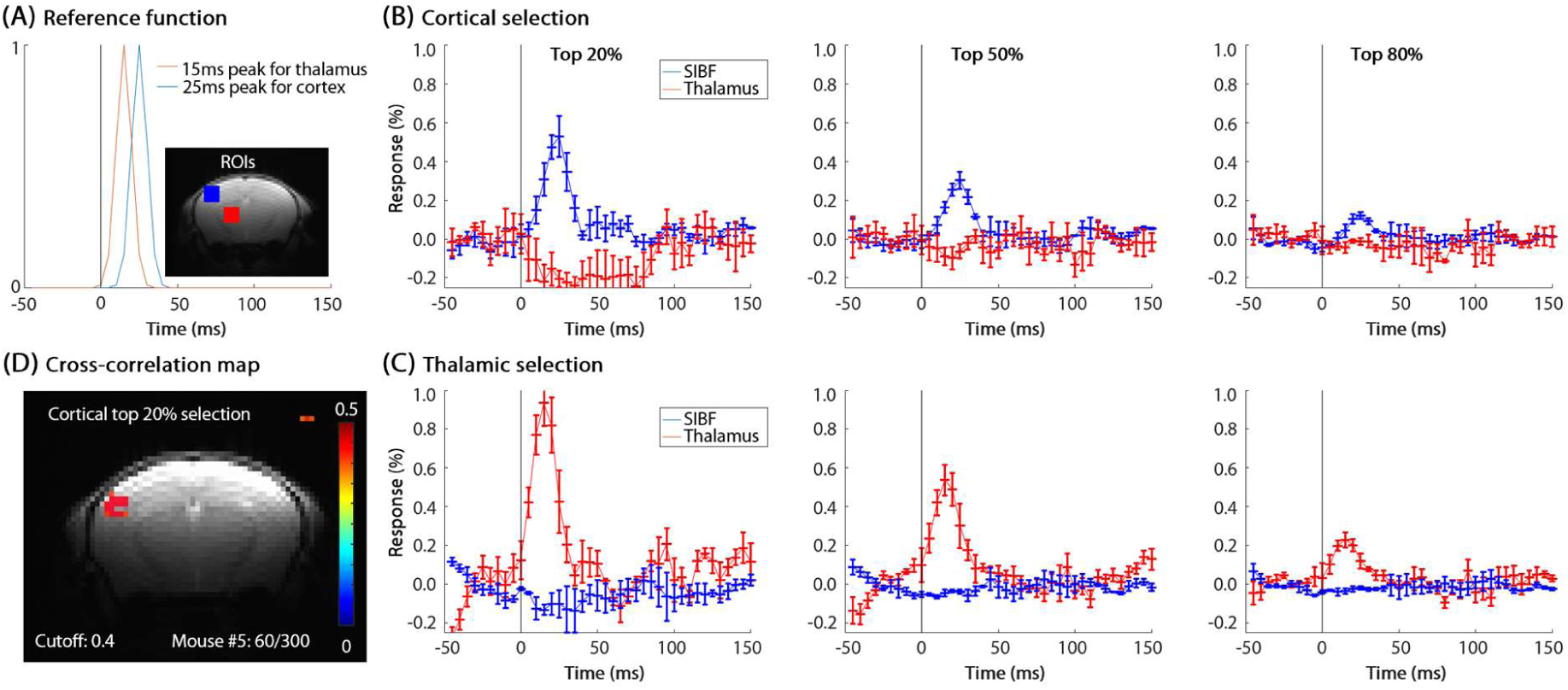
No DIANA-like response in the functionally networked region from trials selected in only one region. (**A**) Gaussian reference functions for 5×5-voxel regions of the active contralateral thalamus (red) and S1BF (blue). Neuronal responses in the thalamus and S1 are expected to peak at ∼15 ms and ∼25 ms after the onset of whisker stimulus (Figure 2 in Toi et al. (*10*)), respectively. In three mice with significant BOLD responses also in the contralateral thalamus, a Gaussian reference function with a FWHM of ∼15 ms peaked at 25 ms in the S1BF (blue) and 15 ms in the thalamus (red) was used to cross-correlate time courses of individual trials. In each animal, trials were selected, based on a threshold of top 20%, 50% and 80% cross-correlation values in each region. Two sets of selected trials (i.e., cortical and thalamic selection) for each threshold were obtained. (**B** and **C**) Averaged time courses of the S1BF and thalamus ROI obtained from the S1BF-(C) and thalamus-selected trials (D) in 3 mice. Blue: S1BF time course; red: thalamus. Artifactual peak intensity in the *selected* region was closely dependent on the selection threshold, while no obvious peak was observed in the *unselected* region. Error bars: SEM. (**D**) Cross-correlation map of one mouse (Mouse #5). For the top 20% trials selected for the S1BF, voxel-wise cross-correlation values with Gaussian neural response functions peaking between 10 ms and 30 ms were calculated, and the highest cross-correlation values were mapped with a correlation amplitude threshold of 0.4 and a minimum of three contiguous voxels (see also fig. S4).

### DIANA observations in the original paper can be replicated from a priori trial selection from noisy data

Our next question is how Toi et al. achieved consistent DIANA responses and maps corresponding to evoked neuronal activity in various experimental conditions (*10*). Despite the lack of description in the original paper (*10*), trial selection was alluded to by the authors (*21*). To examine whether the original DIANA findings can be reproduced using biased signal processing, we selected trials based on the sum of cross-correlation values in both the S1BF and thalamus regions (with reference functions shown in Fig. 5A) in all six mice. The presumable DIANA peak with a ∼10 ms difference in S1BF and thalamus in the original paper (see Figure 2 in Toi et al. (*10*)) was reproduced through the combined selection of trials (Fig. 6A-B). To further investigate whether DIANA maps shown in the original paper (10) can be replicated, we adopted similar processing approaches used in Toi et al. (*10*), including a 3-point moving average and a 3 × 3-voxel median filter, for the top 20% selected trials. In the cross-correlation and time shift maps (Fig. 6C-D for one mouse and fig. S5 for all 6 mice), functional clusters in the S1BF and thalamus were clearly observed in two animals with 60 selected trials (Mouse #5 and #6), replicating DIANA maps in the original paper (see Figure 2 in Toi et al. (*10*)). These indicate that the reported DIANA responses in the original paper (*10*) are likely artifacts resulting from insufficiently or selected averaged data.

**Figure 6.**
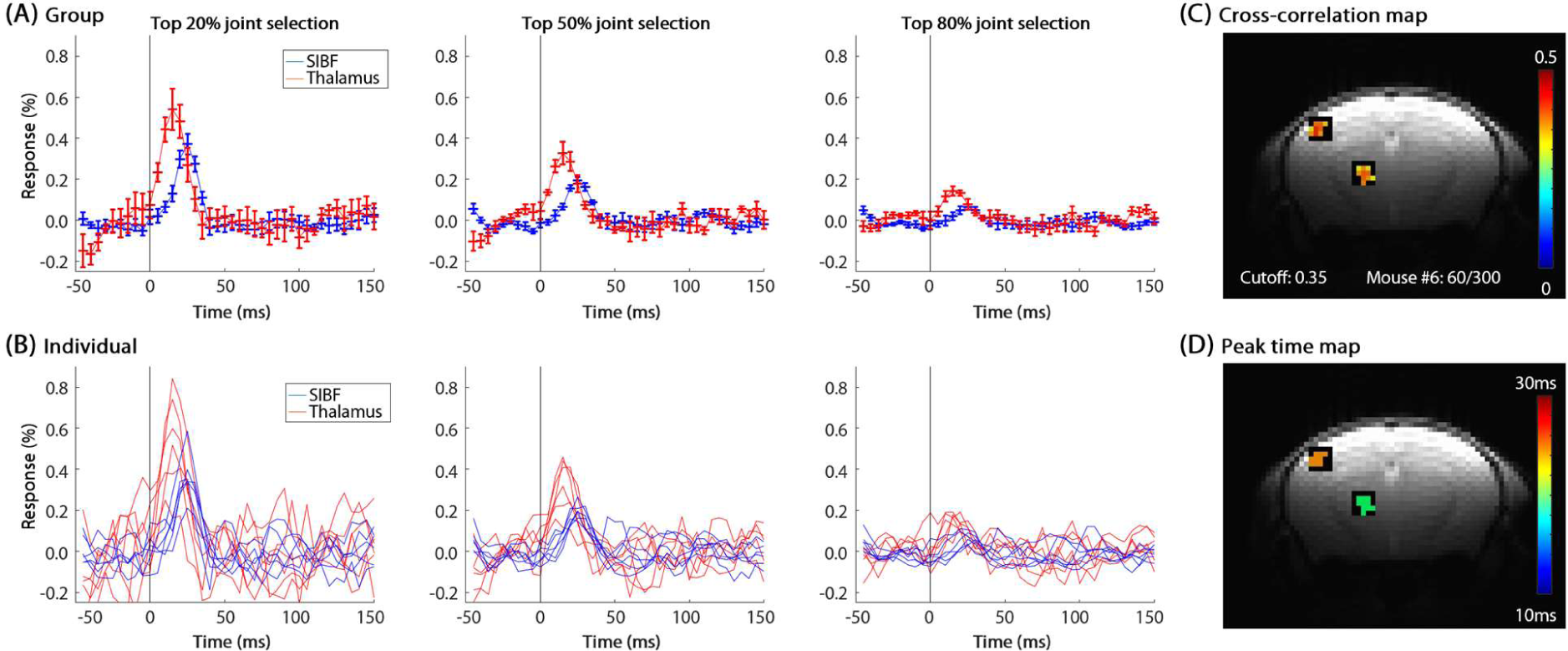
Replication of artifactual DIANA findings in the S1BF and thalamus from trials selected in both regions. A Gaussian reference function with a FWHM of ∼15 ms peaked at 25 ms in the S1BF and 15 ms in the thalamus was used to cross-correlate time courses of individual trials (see Fig. 5A). In all six individual mice, trials were selected jointly from both ROIs, based on a threshold of top 20%, 50% and 80% combined cross-correlation values. **(A-B)** Averaged (A) and individual animal’s time courses (B) of the S1BF and thalamus ROI obtained from the jointly selected trials in 6 mice. Artifactual peaks with a 10 ms difference between thalamus and S1BF were obviously observed as expected. Error bars: SEM. **(C-D)** Cross-correlation (C) and peak time map (D) of one mouse (Mouse #6). For top 20% trials selected for both S1BF and thalamus, voxel-wise cross-correlation values with Gaussian neural response functions peaked between 10 ms and 30 ms were calculated, and the highest cross-correlation values and time shifts were then mapped with a cross-correlation threshold of 0.35 and a minimum of three contiguous voxels (see also fig. S5).

## Discussion

In our high temporal resolution fMRI studies with 50 – 300 averages at 15.2 T, ROI-wise tSNR was ∼2,000 to ∼4,000 in the somatosensory barrel cortex, values that are very high by normal BOLD fMRI standards. A tSNR of 3000 to 1 corresponds to a standard deviation of 0.033%, and thus our data have more than sufficient statistical power to detect the reported ∼0.15% DIANA peak in individual animals. However, we did not find any direct neuronal-related signal in our extensively averaged fMRI studies (Fig. 3), suggesting that the lack of observation of the DIANA peak cannot be due to insufficient tSNR. Since BOLD responses were reliably observed in all the animals, poor control of anesthetic depth is also not a cause of no DIANA observations. Our data suggest that the reported DIANA peak in anesthetized mice (*10*) is not likely neuronal-related fMRI peak, but artifacts due to the noise characteristics of the insufficiently or selected averaged data. This is apparent in Fig. 2B, where spatially distributed random voxels can be found that match the cross-correlation template when using only 50 trials. This random distribution shows apparent activation in the S1BF, the thalamus and many other regions of the brain. This random pattern is in contrast to the BOLD activation maps in Fig. 1C and Fig. 1F.

The exact source of differences between ours and that of Toi et al. (*10*) remains unknown. The lack of replication of the original DIANA findings may be attributed to variations in field strength (15.2 T vs. 9.4 T), surface coil diameter (15 mm vs. 10 mm), and anesthesia protocol (continuous IV infusion vs. bolus IP injection). Firstly, we opted for a higher magnetic field strength of 15.2 T over 9.4 T, despite the availability of both systems, as higher magnetic fields enhance SNR. Concerns about larger BOLD contributions at higher magnetic fields were considered (4,19). However, the minimal BOLD contribution to DIANA signals was expected due to the short TE of 2 ms and the temporal mismatch between fast DIANA and slow BOLD responses (“Suppression of the BOLD signal: Theoretical analysis” section in Supplementary Materials, ref. (10)). The authors of the original paper ascribe the DIANA contrast to T_2_ effects, which should be enhanced at 15.2 T due to the shorter T_2_ of tissue water. Secondly, in our prior field-dependent mouse BOLD fMRI studies (15), we compared results obtained with a 15 mm diameter coil at 15.2 T and a 10 mm diameter surface coil at 9.4 T. Using identical imaging parameters (TE/TR = 16/1000 ms), the average tSNR at 15.2 T vs. 9.4 T was 86 vs. 74 in the forelimb S1 and 63 vs. 45 in the thalamus. This demonstrated that our setup has higher detectability, particularly in the thalamus, compared to the original DIANA studies (10). Thirdly, it should be noted that we initially devised the bolus IP injection protocol (14) for mouse fMRI studies. In this protocol, heart rates decreased and reached a steady-state level within approximately 7 minutes after the bolus injection (refer to Fig. 2 in ref. (14)). The BOLD response was notably weak during the first fMRI session following the initial IP injection but stabilized after the second injection (refer to Fig. 6 in ref. (14)), indicating a deep initial anesthetic depth with variations during repeated bolus injections. To address the challenge of unstable anesthetic depth, we recently introduced an improved continuous infusion protocol (12). Over an 8-hour period following the initiation of ketamine/xylazine IV infusion, heart rate and respiratory rate remained stable (refer to Fig. 1 in ref. (12)), resulting in consistent BOLD responses to whisker pad stimulation (refer to Fig. 2 in ref. (12)). These findings suggest that the continuous infusion protocol is superior to intermittent IP injection for ensuring the reproducibility of repeated fMRI trials. It should be noted that, unlike awake behaving animals, individual whisker pad electric pulses should consistently induce evoked neural activity in the primary somatosensory cortex of lightly anesthetized mice. In our previous studies involving 30-second somatosensory stimulation in rats under isoflurane (22), robust evoked neural activity in response to each pulse was consistently observed, despite a reduction in magnitude with repeated stimuli (refer to Fig. 1 in ref. (22)). Thus, the observed variation in evoked neural activity and fMRI response in anesthetized mice is most likely attributed to the modulation of anesthetic depths (23–25). During the International Society for Magnetic Resonance in Medicine Annual meeting in 2023 (*21*), the senior author of Toi et al. (*10*) commented that averaging more trials in their time series data did not increase tSNR. In our data, tSNR increases with increased number of trials, as expected for stimulus-locked trials with a random noise background (see fig. S1). The difference of variance characteristics between Toi et al. (*10*) and ours can be explained by the stability of anesthesia level-dependent animal physiology during repeated trials. Summed up together, the difference in experimental protocols cannot explain the lack of replication of DIANA peaks in our studies.

When time points and averages (i.e., sample size) are limited, then it is possible that a noise peak can be mistakenly identified as a genuine peak as seen in Fig. 2. This potential problem increases with decreasing tSNR. To separate between genuine peaks and artifacts, extensive averaging should be performed until a peak amplitude is much greater than baseline signal fluctuations (see Fig. 3). Alternatively, if each run has sufficient temporal frames with repeated stimuli, then we can evaluate whether the assumed peak is reproducible across different stimulation pulses. Thus, one should be extremely careful when trying to identify and interpret an unknown peak from limited sampled data.

Our data indicate that a spurious, neuronal activity-like peak can be generated from noisy data with limited time points and trials when some trials are excluded as non-responders by comparison to the hypothesized signal change. We were able to reproduce DIANA signal characteristics observed in Toi et al. (*10*) by the exclusion of outlier-like trials from noisy time courses. For most practical fMRI studies, a few tens of repetitions are all that are acquired due to the limited experimental time *in vivo*. Here we show that the exclusion of even a few trials based on selective filtering may produce artifactually positive findings which are dependent on variances (Fig. 4 vs. fig. S2), but are not seen when no outliers are rejected (Fig. 4 and fig. S2). This demonstrates that positive results can be produced from selective exclusion of spurious time series. Extreme care should be taken to ensure that trial exclusion is based on unbiased metrics such as separately measured BOLD response magnitude or simultaneously acquired physiological parameters and electrophysiology, so that circularity is avoided. Objective, statistically justifiable inclusion/exclusion criteria, having no bias based on any temporal features of the hypothesized result, should be used for data screening and for ensuring that overall findings are preserved, regardless of the exclusion criteria.

While the biased metric approach was the only way that we were able to reproduce the findings of Toi et al. (*10*) (Fig. 6), it is important to emphasize that they did not report using this approach in their paper. In their presentation in the International Society for Magnetic Resonance in Medicine Annual Meeting (*21*), they argued that an increase in the number of repeated trials did not improve DIANA signals due to the contamination of “bad” trials with a certain phase of spontaneous neuronal activity, since the deterministic DIANA response to evoked neuronal activity is modulated by spontaneous ongoing activity. This indicates that the separation of presumably evoked neuronal activity-dominant vs. non-dominant trials (without independently measured electrophysiology) is needed to obtain the DIANA peaks from limited trials, leading to unconfirmed, but possible inclusion of “good” response-appearing trials (Figs. 4-6). To test whether the data selection process is justifiable, we selected “highly neuronal response-like” trials in one area, which were then used to obtain time courses in the functionally networked region (Fig. 5). However, we failed to replicate the findings of the thalamus and S1BF DIANA responses with ∼10 ms time difference in Toi et al. (*10*) from selected trials in only one region, suggesting that the simple trial selection is insufficient to detect the DIANA peak. When we selected trials based on both S1BF and thalamus regions, the DIANA maps and time courses in the original paper (*10*) were artifactually replicated from noisy data (Fig. 6). It should be mentioned that in other measures, such as with Evoked Response Potentials and conventional electrophysiology, also containing both evoked and spontaneous activity, no such selection has ever been reported as necessary to eliminate spurious spontaneous activity, and the statistical power increases, as expected, from more averages.

In conclusion, we could not replicate the original DIANA findings (*10*) even with more extensive averaging, a more precise and steady anesthetic protocol, and higher field strength when no improper data selection was performed. Since subtle differences in pulse sequence or stimulus quality (see Materials and Methods) are unlikely to be substantial, the original DIANA findings are likely to be artifacts due to unreported data processing approaches of noisy data with limited temporal points and trials.

## Materials and Methods

### Animal preparation and stimulation

Six adult male C57BL/6 mice (23-27 g, 10-12 weeks old; Orient Bio, Korea) were used with approval by the Institutional Animal Care and Use Committee (IACUC) of Sungkyunkwan University in accordance with the standards for humane animal care (SKKUIACUC2022-09-55-2). All MRI experiments were performed under anesthesia in accordance with the guidelines of the Animal Welfare Act and the National Institutes of Health Guide for the Care and Use of Laboratory Animals. For anesthesia, a mixture of ketamine and xylazine (100 mg/kg and 10 mg/kg, respectively) was initially injected intraperitoneally (IP), and a dose of ketamine 45 mg/kg/h and xylazine 2.25 mg/kg/h was continuously infused intravenously (IV) (*12, 13, 26*). Note that Toi et al. (*10*) used the supplementary bolus IP delivery of 25 mg/kg ketamine and 1.25 mg/kg xylazine when needed (*9, 14, 15, 27, 28*), which induces anesthetic depth deep right after the bolus injection and a slow recovery to wakefulness (*14, 16*). The animals were self-breathing under continuous supply of oxygen and air gases (1:4 ratio) through a nose cone at a rate of 1 liter/min (*27*). To reduce head motions during image scanning, the mouse was carefully positioned on a customized cradle with two ear plugs, a bite bar and nose cone. Body temperature was maintained at 37 ± 0.5 C^ͦ^ with a warm-water heating system via rectal temperature probe.

For whisker electric pad stimulation, two anodes with 2 mm apart and a cathode (*12*) were placed on the mouse’s right whisker pad. Note that Toi et al. (*10*) used 2×2 anodes which was originally developed for rat whisker pad stimulation (*29*). Pulse parameters were at a pulse width of 0.5 ms, a frequency of 4 Hz (for BOLD studies), and current intensity of 0.5 mA, controlled by a pulse generator (Master 9; World Precision Instruments, Sarasota, FL, USA).

### fMRI data collection

Data were acquired on a 15.2 T (Bruker BioSpec MRI, Billerica, MA, USA) equipped with a 11 cm horizontal bore magnet and actively shielded 6-cm gradient. A 15 mm ID surface coil was used for both transmission and reception. Note that Toi et al. (*10*) used a 10 mm diameter surface coil at 9.4 T, and its SNR and tSNR are lower than those of 15.2 T with 15 mm diameter coil (*15*). Mouse brain was placed at the isocenter of magnet and field inhomogeneity was minimized via MAPSHIM protocol in Paravision 6.0.1 software. Detailed experimental procedures were described in our previous publications (*9, 15*). The 2-D shuffled line scanning pulse sequence (*6, 10, 30*) was modified from the conventional FLASH sequence provided by the vendor. Onset of the stimulation pulse(s) was synchronized with the onset of RF pulse in the first image for each stimulation block.

Three fMRI approaches were used as follows.

1. BOLD-EPI: Standard gradient-echo echo planar imaging (EPI) was used to obtain BOLD fMRI with the following imaging parameters; image matrix = 96 × 48, FOV = 16 × 12 mm^2^ (0.17 × 0.25 mm^2^ in-plane resolution), 1.0 mm-thick slices, TR/TE = 1000/11.5 ms, bandwidth = 300,000 Hz, and 50° flip angle. Then, a single 1-mm slice was chosen based on scout BOLD fMRI (approximately −1.75 mm from bregma) for subsequent fMRI studies. Single-slice BOLD fMRI was performed for ensuring reliable BOLD activity. The stimulation paradigm consisted of 40 s control, 20 s stimulation, 60 s control, 20 s stimulation, and 60 s recovery, lasting 200 s. Two runs of BOLD fMRI were repeated before and after DIANA scans to ensure their stable responsiveness during entire fMRI studies.
2. BOLD line scanning (BOLD-LS): The 2-D shuffled line scanning method (*6, 10, 30*) was adopted for single-slice BOLD fMRI with TR/TE = 100/11.7 ms, FOV =16 × 12 mm^2^, image matrix =72 × 54, bandwidth = 50,000 Hz, slice thickness = 1 mm, and flip angle = 17° in the S1BF (Ernst flip angle with T_1_ of 2.2 s). The stimulation paradigm consisted of 2 s control, 1 s stimulation, 6 s control, 1 s stimulation, and 6 s recovery. Each trial lasted 16 s × 54 phase encoding steps = 864 s. Only one trial was obtained for ensuring correct implementation of the pulse sequence in our MRI system.
3. DIANA: DIANA experiments were performed with the 2-D shuffled line scanning approach with the same imaging parameters used in Toi et al. (*10*), TR/TE = 5/2 ms, 4° flip angle in the S1BF, FOV = 16 × 12 mm^2^, image matrix =72 × 54, slice thickness = 1 mm, and bandwidth = 50,000 Hz. In some studies, RF spoiling and steady-state dummy scans were used (see table S1). Two different numbers of frames for each phase-encoding step were acquired, 40 (200 ms) or 200 frames (1000 ms). For DIANA studies with total 40 frames, 50 ms pre-stimulation (10 frames) and 150 ms post-stimulation images (30 frames) were acquired, lasting 200 ms × 54 phase encoding steps = 10.8 s/trial, which is the same as Toi et al. (*10*) used. Alternatively, 20 pre-stimulus and 180 post-stimulus images, or 20 pre-stimulus and 2 times 90 post-stimulus images were obtained (see table S1). At least 50 trials were repeated.

### fMRI data processing

We did not exclude any dataset, and all the acquired data were analyzed with AFNI software (https://afni.nimh.nih.gov/) (*31*), FMRIB Software (FSL, https://fsl.fmrib.ox.ac.uk/fsl/fslwiki/) (*32*), and Matlab codes (Mathworks).

#### Generating fMRI maps

Each individual animal’s fMRI maps were generated using preprocessing and a cross-correlation analysis with a hemodynamic response function (HRF) (*15*) or an expected DIANA response function (*10*). The DIANA response function was modeled as a Gaussian window function in Matlab, ‘gausswin’, with a peak position at 25 ms after the stimulus and the full width at half maximum (FWHM) of ∼25 ms. The following preprocessing steps were performed to improve the detection of signal activation: linear detrending for signal drift removal and normalization by the average of the pre-stimulus baseline volumes. All repeated fMRI trials on each animal were averaged and spatial smoothing was applied using a Gaussian kernel with FWHM of one voxel.

#### ROI analysis

Active and inactive ROI with 5×5 voxels was chosen in the contralateral S1BF and the ipsilateral thalamic area from BOLD-EPI functional maps, respectively. In addition, the contralateral thalamus ROI with 5×5 voxels was selected based on activation maps (see Fig. 1C). Since EPI and line scanning used a slightly different spatial resolution, the ROIs determined from BOLD-EPI were similarly positioned in the line scanning images. In each run (BOLD-EPI, BOLD-LS, DIANA), signals in the selected 5×5-voxel ROI were averaged from detrended, normalized image frames (without any spatial or temporal smoothing), then were temporally smoothed with 3-point moving averages, which was used in Toi et al. (*10*). In each subject, repeated DIANA runs with 10 pre-stimulus and 30 post-stimulus frames were temporally aligned, based on the stimulus time. The 10^th^ image (right before the stimulus pulse) was set to 0 ms for being consistency with Toi et al. (*10*). Since TE of 2 ms was used, the exact time of the 11^th^ image (the image right after the stimulus pulse) was 2 ms, but assigned to 5 ms. The percent signal change time course was calculated by 100×ΔS/S_base_, where ΔS is a difference between signals at the reference and at the baseline period and S_base_ is the averaged baseline pre-stimulus signal (1^st^ to 10^th^ frame). In each subject, 50 – 300 trials were averaged. Finally, averaged subject-wise time courses were averaged. Since different animals had different number of averages, a grand total average was also performed from 1050 trials of the six subjects.

#### BOLD percent change and tSNR calculation

In each subject, ROI-wise BOLD-EPI responses to whisker stimulation were calculated from 0 – 35 s time frames after the stimulus onset, while ROI-wise BOLD-LS changes were calculated from 1 – 4.5 s time frames after the stimulus onset. To determine the detectability of DIANA measurements, tSNR values were calculated by means divided by its standard deviations from time series data of individual voxels or ROIs. Although tSNR calculation generally uses only pre-stimulus data points, we used the entire time series due to no obvious observation of stimulation-induced responses.

#### Histogram analysis

Histograms of individual DIANA time series were obtained by counting the number of data points every 0.1 bins. A Gaussian fitting to histogram was performed and its mean and standard deviation were determined.

#### Selection of trials

To evaluate the impact of trial exclusion (Fig. 4), each trial-wise ROI time course with 3-point moving averaging was cross-correlated with a hypothetical outlier Gaussian function with a negative peak time of 25 ms as an expected neuronal time (and 35 ms as a control) and FWHM of ∼25 ms. Then, trials were ranked, based on cross-correlation values. Various thresholds of top 6%, 10%, and 20% were used for the separation between included and excluded trials. Then, the included and excluded trials were separately averaged for each animal (Fig. 4).

To determine the validity of trial selection (Fig. 5), each trial-wise ROI time course (with 3-point moving average) of contralateral S1BF (thalamus) was cross-correlated with a hypothetical Gaussian function with a positive peak time of 25 ms (15 ms) and FWHM of ∼15 ms. Then, top 20%, 40%, and 80% threshold trials were selected separately for S1BF and thalamus ROI. Then, time courses of S1BF and thalamus were obtained from each selected set of trials for S1BF and thalamus (Fig. 5 and fig. S3). For the top 20% trials selected for S1BF, we applied one-voxel spatial smoothing and a 3-point moving average. Subsequently, an averaged trial was cross-correlated with Gaussian neural response functions peaking between 10 ms and 30 ms on a voxel-by-voxel basis. The highest cross-correlation values and their corresponding peak times were then determined (Fig. 5 and fig. S4).

In addition, trials were selected by the threshold of top 20%, 40%, and 80% of *combined* cross-correlation values for the thalamus and S1BF reference functions (see Fig. 6). Then, time courses of S1BF and thalamus were obtained from the *commonly selected* trials (Fig. 6). For the top 20% trials selected for the combined S1BF and thalamus, 3-point moving averaging was performed. Then, an averaged trial was cross-correlated with Gaussian neural response functions peaking between 10 ms and 30 ms on a voxel-by-voxel basis. The highest cross-correlation values and their peak times were then determined. Following that, a 3 × 3-voxel median filter was applied to obtain DIANA fMRI maps, and the highest cross-correlation values along with their peak times were presented (Fig. 6 and fig. S5).

#### Statistical analysis

All statistical analyses between the two BOLD groups (Fig. 1) were conducted with paired t-test. To determine a statistical significance in Fig. 2, all pre-stimulus baseline data (20 time points × 50 trials) were compared with peak data points (1 time point × 50 trials) with two-group t-test, which was similarly done in Toi et al. (*10*). Quantitative values were presented as mean ± standard deviation and plots were presented as mean ± standard error of the mean (SEM).

## Acknowledgments

We thank Prof. Jung Hee Lee for valuable discussion.

## Funding

This research was supported by the Institute of Basic Science (IBS-R015-D1 to S.-G.K.), and by NIH (RF1NS113278, R01NS124778, R01NS122904 to X.Y and S.C).

## Author contributions

Conceptualization: SGK, XY

Methodology: SC, XY

Investigation: SHC, GHI, PAB, RSM

Visualization: SHC

Supervision: SGK

Writing—original draft: SGK

Writing—review & editing: SGK, SHC, PAB, RSM, SC

## Competing interests

The authors declare no other competing interests.

## Data and materials availability

All data needed to evaluate the conclusions in the paper are present in the paper and/or the Supplementary Materials. The unprocessed fMRI data are available from the corresponding author upon request.

## Supplementary Materials

Figs. S1 to S5

Table S1

## Notes

### Competing Interest Statement

The authors have declared no competing interest.

### Summary of Updates

The content of the manuscript has been revised and expanded overall, including the abstract and supplementary material.

