## Supplementary_materials for "No Replication of Direct Neuronal Activity-related (DIANA) fMRI in Anesthetized Mice": Supplementary_materials_noDIANA.pdf


**The PDF file includes:**

Figs. S1 to S5
Table S1

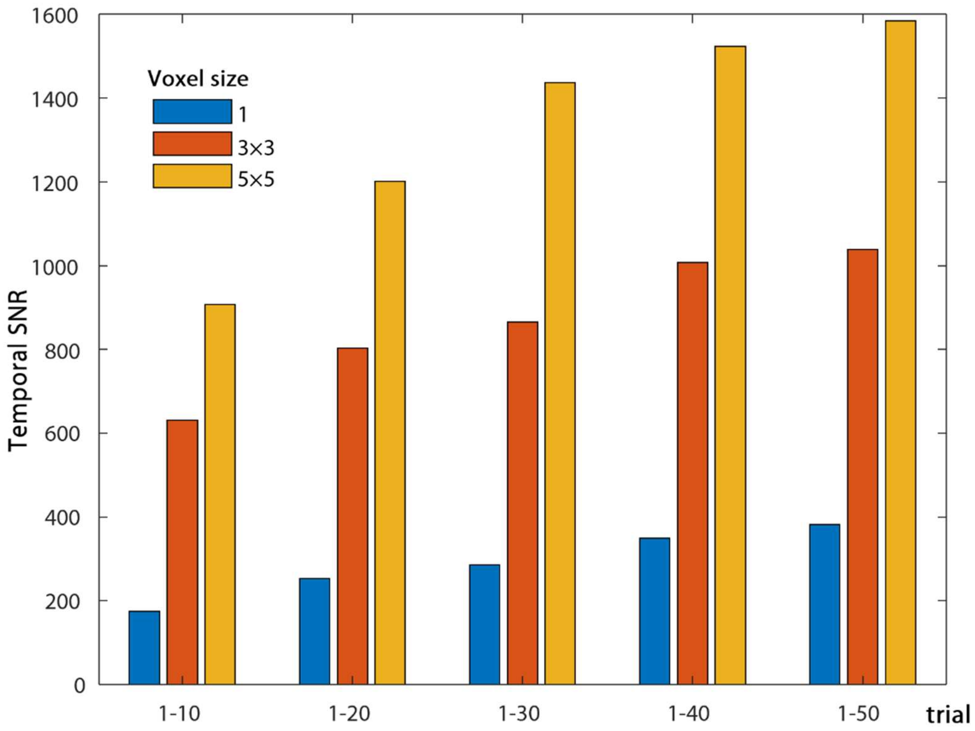


**Fig. S1. Systematic analysis of temporal signal-to-noise ratios (tSNR) within the S1BF ROI as a function of averages and voxel numbers**. tSNR was obtained from 200 frames with TR of 5 ms from 50 DIANA trials in one animal (Mouse #3, see table S1). One center voxel, and middle 3×3 voxels, and all 5×5 voxels were chosen within the 5×5-voxel S1BF ROI. To examine the effect of averages, the first 10, 20, 30, 40, and all 50 trials were averaged. Clearly, tSNR increases with spatial and temporal averages as expected by random noise theory.

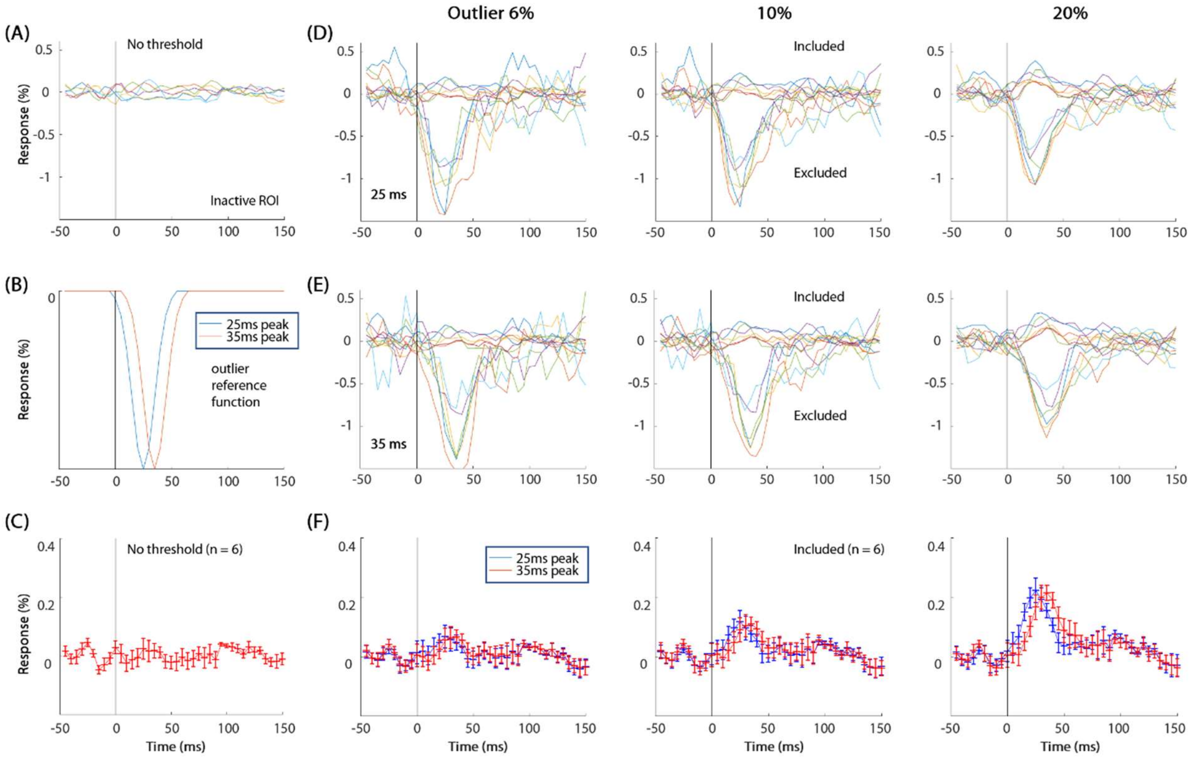


**Fig. S2. Effect of temporal response-based data exclusion in the inactive ROI.** (**A**) Averaged time courses of six individual animals in the inactive ROI without excluding trials (same as Fig. 3D). (**B**) A Gaussian reference outlier function with FWHM of ~25 ms peaked at 25 ms (blue) and 35 ms (red). (**C**) An average of all animals' time courses without excluding trials. This is the same as Fig. 3H. In six animals, an individual trial's time course was correlated with a reference function shown in B, and ranked among all trials, based on its cross-correlation value. Then, in each animal, trials were separated into included and excluded categories, based on an outlier threshold of top 6%, 10%, and 20% cross-correlation values. (**D** and **E**) Averaged time courses of the included and excluded trials in individual animals for the 25 ms (**D**) and 35 ms peak reference function (**E**). Averaged time courses of the included trials showed noisy positive responses, while those of the excluded trials showed a negative peak around the expected peak time. Each color time course: each animal. (**F**) Means of subject-wise included trials time courses for 25 ms (blue) and 35 ms peak reference functions (red). Clearly, an exclusion process leads to an erroneous peak from noisy data. It is fundamentally important to note that these peaks are erroneous because such a preselection process is statistically circular, summing noise in a biased manner to produce precisely the peak that was preselected. Error bars: SEM.

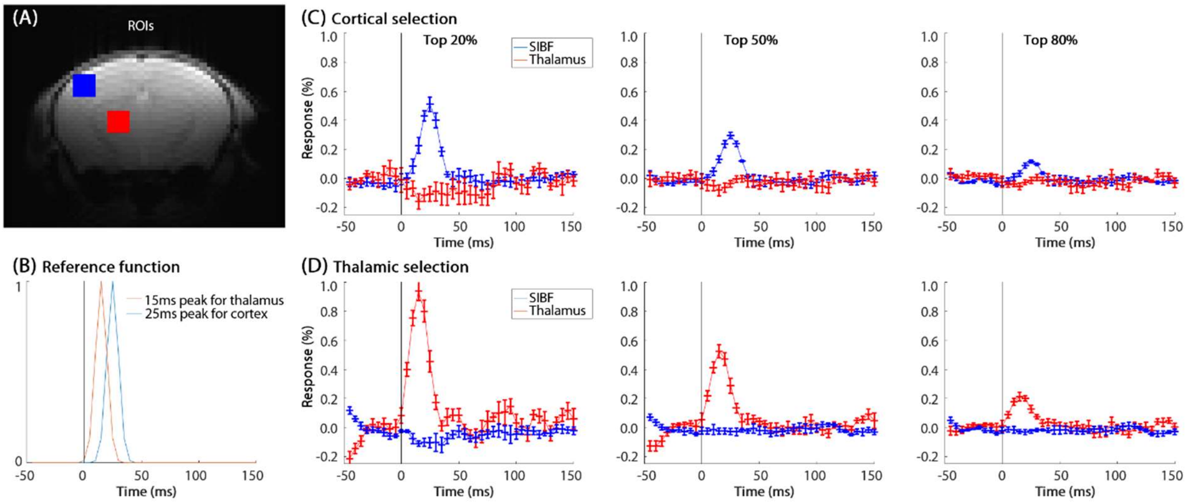


**Fig. S3. No DIANA-like response in the networked somatosensory region in all 6 mice from trials selected in only one region.** (**A**) Regions of the contralateral thalamus and S1BF. Neuronal responses in the thalamus and S1 are expected to peak at ~15 ms and ~25 ms after the onset of whisker stimulus (Figure 2 in Toi et al. (10)). (**B**) A Gaussian reference function with a FWHM of ~15 ms peaked at 25 ms in the S1BF (blue) and 15 ms in the thalamus (red) was used to cross-correlate time courses of individual trials. In each animal, selected trials were averaged, based on a threshold of top 20%, 50% and 80% cross-correlation values. (**C** and **D**) Averaged time courses of the S1BF and thalamus ROI obtained from the S1BF- (C) and thalamus-selected trials (D) in 6 mice. Artifactual peak intensity in the *selected* region was closely dependent on the selection threshold, while no obvious peak was observed in the *unselected* region. Error bars: SEM.

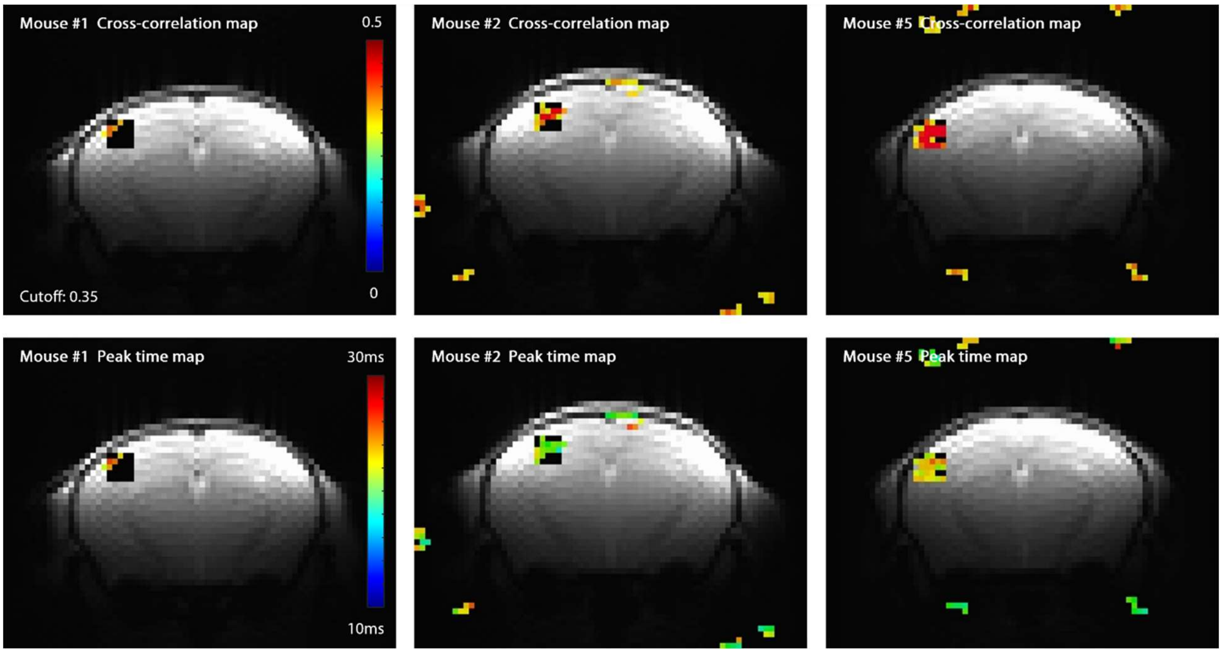


**Fig. S4. Cross-correlation (top row) and peak time map (bottom row) of the top 20% trials selected for the S1BF in the thalamus-activated 3 mice (Mouse #1, #2, and #5).** The mapping procedure was identical to that of Fig. 5D. For the top 20% of trials selected for the S1BF, voxel-wise cross-correlation values were calculated with Gaussian neural response functions peaking between 10 ms and 30 ms. The highest cross-correlation values and their peak times were then mapped with a correlation amplitude threshold of 0.35 and a minimum of five contiguous voxels. The black square region in each image indicates the S1BF ROI. Only the S1BF cluster was active, as expected by circular reasoning.

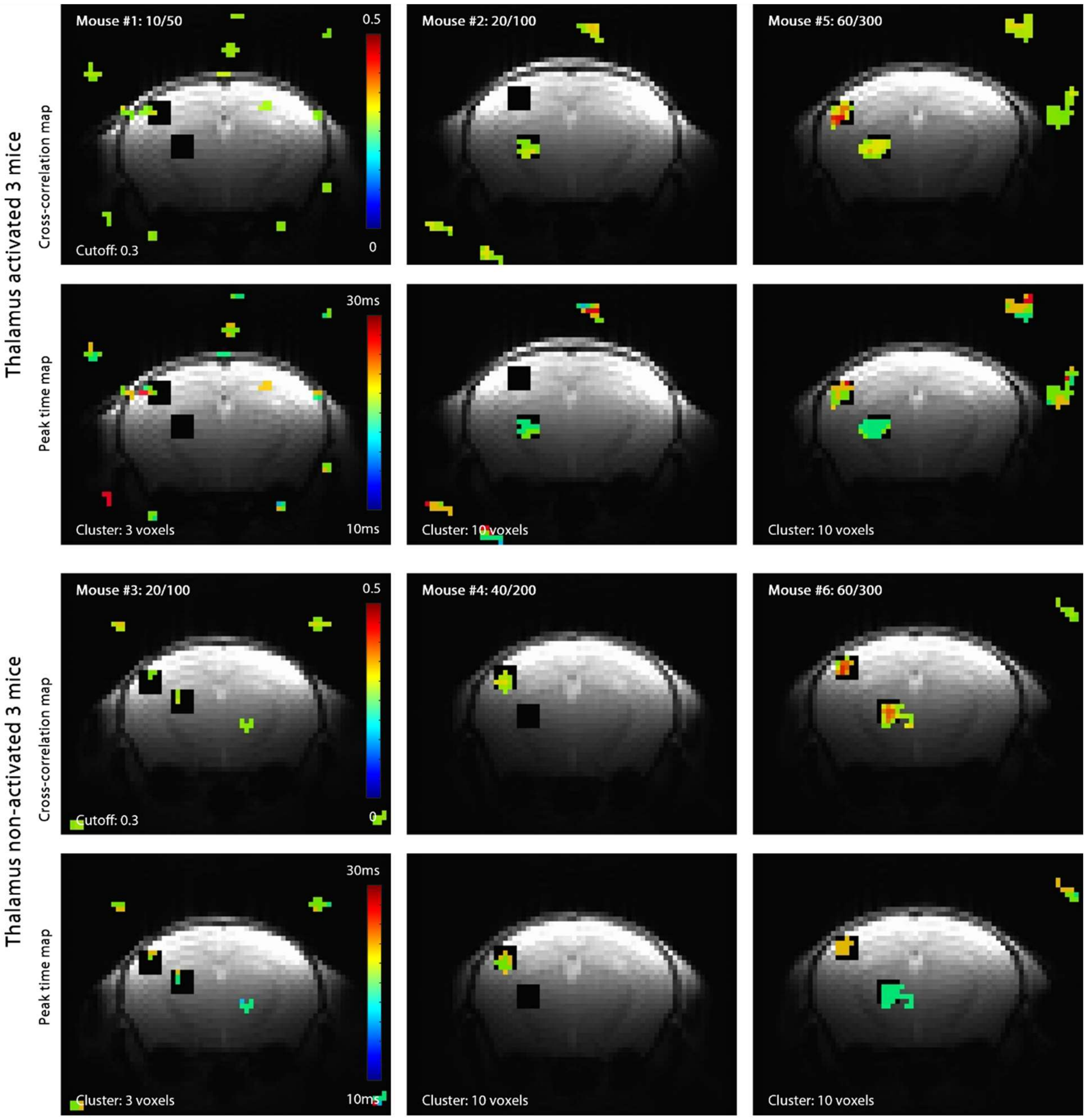


**Fig. S5. Replication of DIANA maps calculated from selected trials in both regions for all 6 mice.** The upper panel displays three mice with significant BOLD activation in the thalamus, while the bottom panel shows three mice without significant BOLD activation in the thalamus. The mapping procedure was identical to Fig. 6C-D. For the top 20% of trials selected for combined S1BF and thalamus, voxel-wise cross-correlation values were calculated with Gaussian neural response functions peaking between 10 ms and 30 ms. Maps for the highest cross-correlation values (first row) and their time shifts (second row) were obtained with a cross-correlation threshold of 0.3 and a minimum of either 3 or 10 contiguous voxels. The upper-left corner of the cross-correlation map indicates the number of selected trials out of the total number of trials. Two black square regions in each image indicate the S1BF and thalamus ROIs. When 60 trials were selected (Mouse #5 and #6), DIANA maps in the original paper were successfully replicated by improper signal processing of noisy data.

**Table S1. Detailed experimental parameters.**

| Mouse ID | Paradigm[*] (ms) | ISI[†] (s) | Pulse # | Average | Trial #[‡] | RF-spoil[§] | Steady State |
|---|---|---|---|---|---|---|---|
| 1 | 50–150 | 0.2 | 1 | 50 | 50 | x | x |
| 2 | 50–150 | 0.2 | 1 | 100 | 100 | x | x |
| 3 | 50–150 | 0.2 | 1 | 50 | 100 | x | x |
| | 100–900 | 1.0 | 1 | 50 | | x | x |
| 4 | 50–150 | 0.2 | 1 | 100 | 200 | x | x |
| | 100–450–450 | 1.0 | 2 | 50 | | x | x |
| 5 | 50–150 | 0.2 | 1 | 100 | 300 | x | o |
| | 50–150 | 0.2 | 1 | 100 | | o | o |
| | 100–450–450 | 1.0 | 2 | 50 | | o | x |
| 6 | 50–150 | 0.2 | 1 | 100 | 300 | x | o |
| | 50–150 | 0.2 | 1 | 100 | | o | o |
| | 100–450–450 | 1.0 | 2 | 50 | | o | x |
| Total | 50–150 | | 1 | | 1050 | | |

[*]Stimulation paradigm. "–" indicates a 0.5-ms single whisker pad electric stimulus, and image TR is 5 ms.
[†]Inter-scan interval.
[‡]Number of trials time-locked with a stimulus pulse.
[§]Radio frequency (RF) spoiling.
